# Temporal control of acute protein aggregate turnover by UBE3C and NRF1-dependent proteasomal pathways

**DOI:** 10.1101/2024.08.30.610524

**Authors:** Kelsey L. Hickey, Alexandra Panov, Enya Miguel Whelan, Tillman Schäfer, Arda Mizrak, Ron R. Kopito, Wolfgang Baumeister, Rubén Fernández-Busnadiego, J. Wade Harper

## Abstract

A hallmark of neurodegenerative diseases is the progressive loss of proteostasis, leading to the accumulation of misfolded proteins or protein aggregates, with subsequent cytotoxicity. To combat this toxicity, cells have evolved degradation pathways (ubiquitin-proteasome system and autophagy) that detect and degrade misfolded proteins. However, studying the underlying cellular pathways and mechanisms has remained a challenge, as formation of many types of protein aggregates is asynchronous, with individual cells displaying distinct kinetics, thereby hindering rigorous time-course studies. Here, we merge a kinetically tractable and synchronous agDD-GFP system for aggregate formation with targeted gene knockdowns, to uncover degradation mechanisms used in response to acute aggregate formation. We find that agDD-GFP forms amorphous aggregates by cryo-electron tomography at both early and late stages of aggregate formation. Aggregate turnover occurs in a proteasome-dependent mechanism in a manner that is dictated by cellular aggregate burden, with no evidence of the involvement of autophagy. Lower levels of misfolded agDD-GFP, enriched in oligomers, utilizes UBE3C-dependent proteasomal degradation in a pathway that is independent of RPN13 ubiquitylation by UBE3C. Higher aggregate burden activates the NRF1 transcription factor to increase proteasome subunit transcription, and subsequent degradation capacity of cells. Loss or gain of NRF1 function alters the turnover of agDD-GFP under conditions of high aggregate burden. Together, these results define the role of UBE3C in degradation of this class of misfolded aggregation-prone proteins and reveals a role for NRF1 in proteostasis control in response to widespread protein aggregation.

## Introduction

A hallmark of neurodegenerative diseases (NDs) is the progressive loss of proteostasis, leading to the accumulation of aberrant, misfolded and aggregated proteins that result in cytotoxicity (1). Although different NDs involve unique proteins such as amyloid-β (Aβ) and tau in Alzheimer’s disease, and α-synuclein in Parkinson’s disease, the general feature of protein misfolding resulting in higher order structures (aggregates) is shared (2, 3). To maintain proteome stability, cells have evolved quality control (QC) pathways that, when necessary, refold (chaperones), or degrade (ubiquitin-proteasome system and autophagy) misfolded proteins (1, 4, 5). Loss of factors involved in these QC pathways, as well as mutations in proteins that promote off-pathway folding intermediates, have been linked to neurodegenerative phenotypes, highlighting the importance of preventing protein aggregates as a mechanism to maintain proteostasis (1, 6, 7). However, studying the role of degradation pathways has remained a challenge, as aggregate formation is typically asynchronous and heterogeneous at the cellular level. As a result, much of the work in the field has examined mechanisms acting on only late-stage aggregates, well beyond the time where quality control systems are able to effectively remove such structures (8–11). Identifying critical quality control pathways is important for the development of quality control-based therapeutic strategies that function during initial phases of protein aggregation.

Here we merge a kinetically tractable and synchronous agDD-based aggregate formation system (12, 13) with selective gene knockdowns and single-cell quantification of agDD-GFP abundance by flow cytometry to identify strategies that mammalian cells employ to degrade aggregation-prone assemblies at defined stages along an aggregation trajectory, thereby preventing the toxic effects of protein aggregates on cellular homeostasis. We found that aggregate abundance and size inversely correlate with their ability to be removed from cells principally through the ubiquitin-proteasome system. With lower misfolding burden, misfolded proteins are degraded in a manner that relies on the HECT-domain ubiquitin ligase UBE3C, and subsequent proteasomal degradation over the course of several hours. When misfolding burden is high, UBE3C alone is not sufficient for agDD-GFP degradation, leading to larger aggregate formation and activation of the transcriptional regulator NRF1 (also called NFE2L1). NRF1 contains a single-pass transmembrane segment, which adopts a type II membrane protein topology within the endoplasmic reticulum (ER) under basal conditions (14–16). In this context, NRF1’s C-terminal DNA-binding and transactivation domains are localized within the ER lumen and are glycosylated. In response to agents that reduce proteasome activity (e.g. inhibition with protease inhibitors such as bortezomib, BTZ), NRF1’s C-terminal region is retrotranslocated into the cytosol in a manner that depends on the AAA-ATPase VCP (also called p97), deglycosylated by cytosolic deglycosylases and the DNA-binding/transactivation region released from the ER tether by the DDI2 endoprotease (15–19). Liberated and deglycosylated NRF1 translocates to the nucleus, where it can promote expression of genes encoding proteasome subunits. Using genetics and cell biologic assays, we find that NRF1 is stabilized during widespread protein aggregation, activating transcription of proteasome subunits. Positively or negatively altering the activity of the NRF1 pathway increases or decreases the cell’s ability to clear protein aggregates particularly under the conditions of high aggregate load. Together, these results provide insight into how cells acutely respond to protein aggregation to promote aggregate clearance.

## Results

### agDD forms amorphous structures by cryo-ET

To overcome the limitations of slow forming aggregation systems used to examine proteostasis responses, we have leveraged a minimally perturbative and temporally controllable protein aggregation system (12, 13). This system features a destabilizing domain (agDD) derived from FK506 binding protein 12 (FKBP12) fused to the fluorescent GFP protein, the stable folding of which relies on association with a synthetic ligand, Shield-1 (S1) (12, 13). In the presence of S1, agDD is properly folded and exhibits diffuse cytoplasmic localization, but upon ‘washout’ of the S1 ligand, the agDD fusion protein immediately (in second to minutes) unfolds and assembles into structures that coalesce into much larger aggregates over time (12, 13). Previous studies have examined the earliest stages of aggregate formation (12, 13) but how aggregates formed in this system are acted upon by cellular quality control systems remains poorly understood.

As an initial approach to examine agDD aggregates, we performed correlative cryo-electron tomography (cryo-ET) at early (10 min) and late (6 h) times after removal of the S1 ligand in HEK293 cells overexpressing agDD (13). Previous studies using cryo-ET have resolved multiple types of aggregate structures, ranging from fibrils (α-synuclein) (10) to ribbons (poly-Gly-Ala derived from C9ORF72 expansion) (20), with Poly-Q repeats derived from Htt forming both amorphous and fibrillar structures (21, 22). At early times during aggregation, we detected amorphous GFP-positive structures (∼500 nm in diameter) that were often adjacent to membranous structures including ER (**Fig. 1A and SI Appendix, Fig. S1A-C, Video S1**). At longer times (6 h), much larger aggregates were visualized (1-2 microns), and in some instances, aggregate material appeared to be incorporated into membranous organelles (**Fig. 1B and SI Appendix, Fig. S1D-F, Video S2**). Large macromolecular assemblies such as ribosomes or proteasomes appeared to be excluded from aggregates, analogous to previous findings with Htt-related Poly-Q aggregates (21) and in contrast with poly-GA ribbons that sequester proteasomes (20). Thus, our cryo-ET data indicate that agDD-GFP aggregates are largely amorphous and closely interact with various cellular structures.

**Fig 1:**
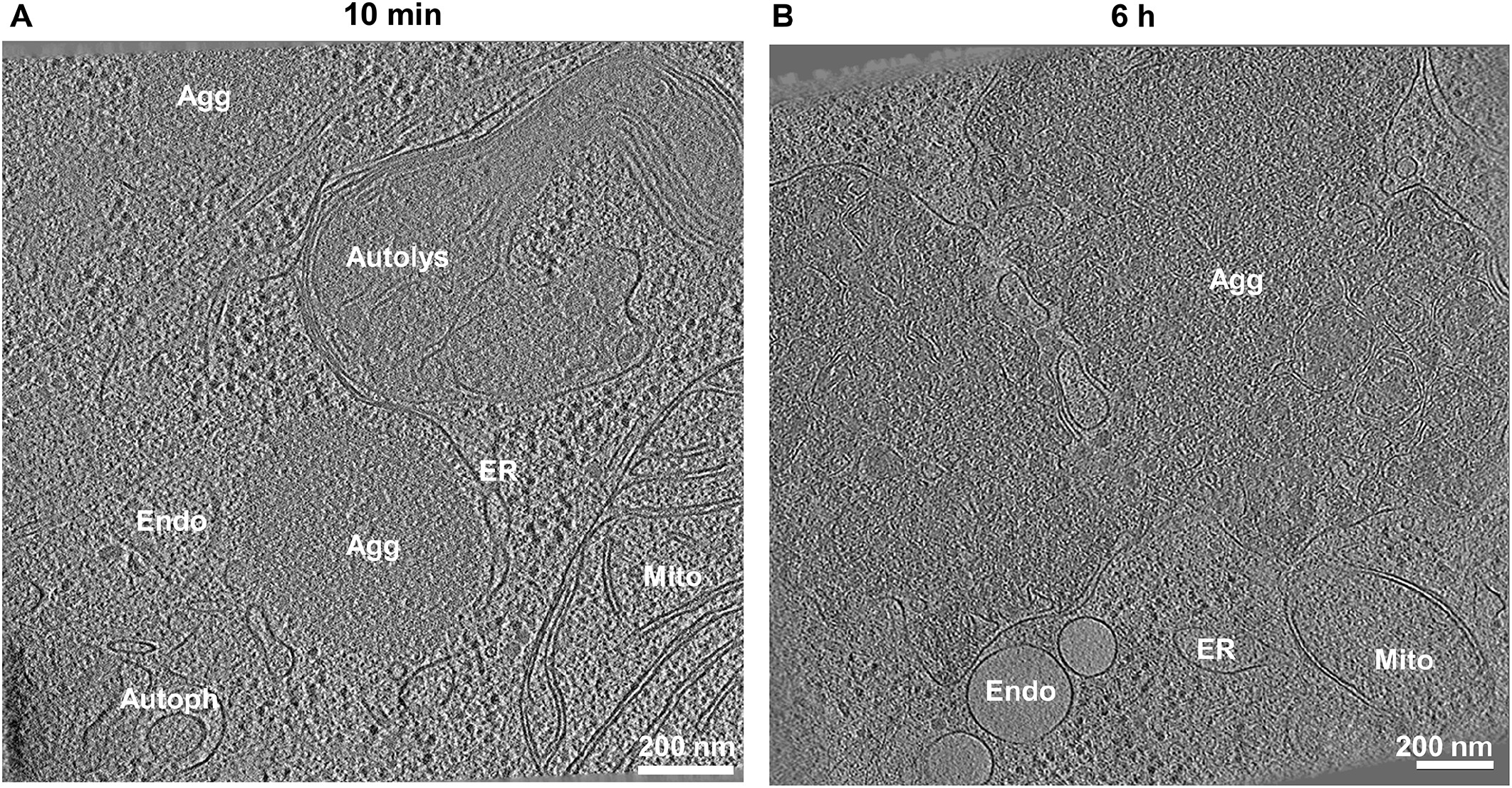
agDD-GFP forms amorphous aggregate structures. **(A)** Tomogram of HEK293 agDD-GFP cells 10 min after S1 washout. **(B)** Tomogram of HEK293 agDD-GFP cells 6 h after S1 washout. Selected cellular structures are annotated. Scale bar, 200 nm. Agg, Aggregate; ER, endoplasmic reticulum; Endo, endosome; Mito, mitochondria; Autolys, autolysosome; Autoph, autophagosome.

### Temporal turnover of agDD is autophagy-independent but proteasome-dependent

Previous studies have examined the formation of agDD-GFP aggregates but the fate of such aggregates has not been examined in detail. HEK293T agDD-GFP cells were either left in S1 or subjected to S1 washout for 4 or 24 h and cells analyzed by flow cytometry to determine GFP fluorescence on a single cell level (**Fig. 2A**). agDD-GFP fluorescence 4h after washout was comparable to cells in the presence of S1, but 24 h post-washout, the mean fluorescence had decreased by ∼10-fold, albeit with a population of cells displaying fluorescence similar to that of control cells persisted (**Fig. 2A**). Given previous data implicating autophagy in turnover of specific types of protein aggregates, we asked whether the reduction in agDD-GFP fluorescence required the core autophagy gene FIP200 (also called RB1CC1), which is considered to provide a full block to canonical autophagy (23). We found, however, that the pattern of agDD-GFP fluorescence seen in WT cells was recapitulated in FIP200^-/-^ HEK293T cells (23) expressing comparable levels of agDD-GFP (**Fig. 2A**, right panel). Thus, canonical autophagy appears to play little or no role in agDD-GFP turnover under these conditions.

**Fig 2:**
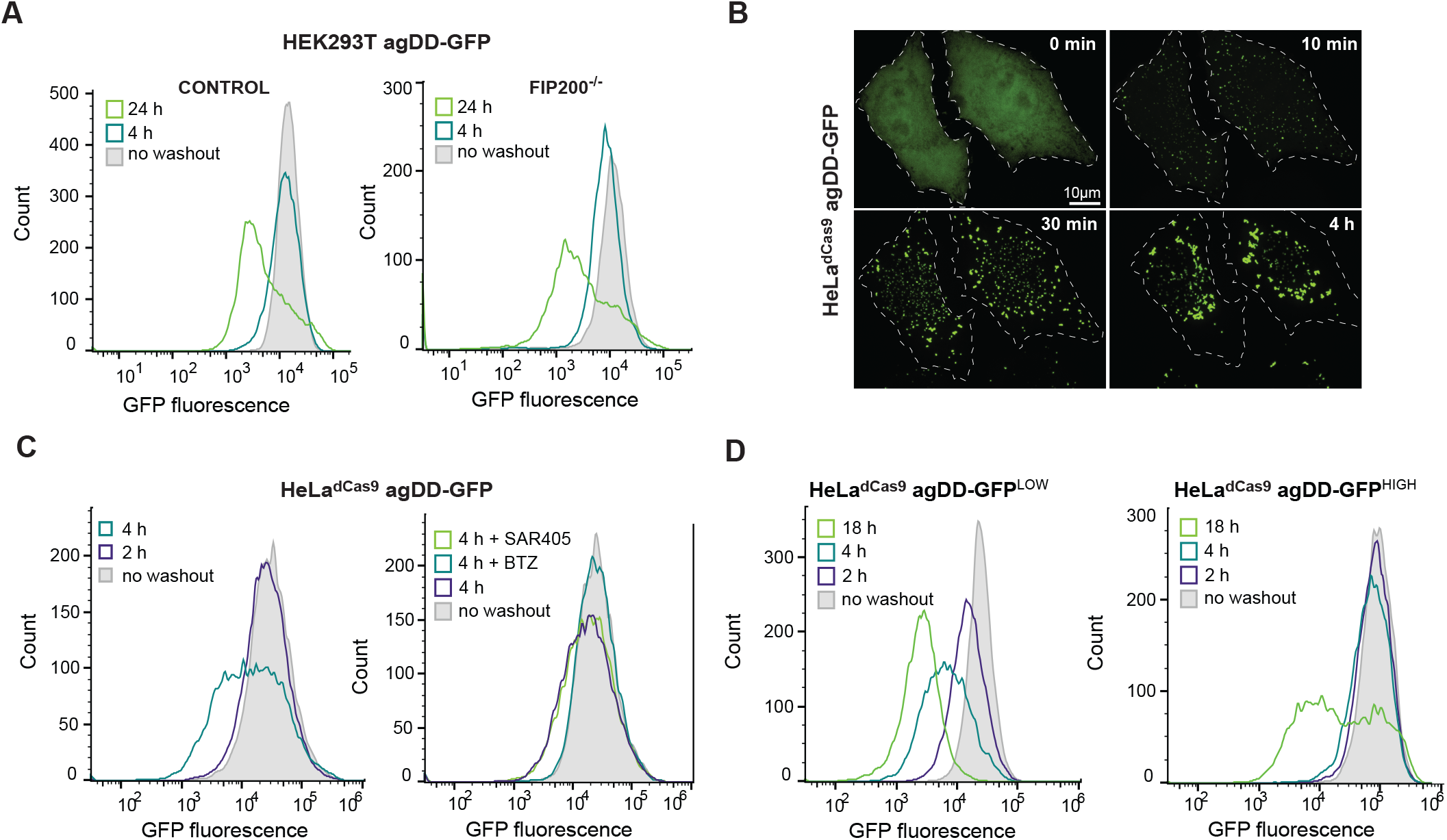
agDD-GFP aggregates are degraded by the ubiquitin-proteasome system. **(A)** Histograms of agDD-GFP fluorescence levels in HEK293T and HEK293T FIP200^-/-^ cells post-S1 washout measured by flow cytometry. **(B)** Live cell confocal imaging of HeLa^dCas9^ agDD-GFP cell line post-S1 washout. **(C)** Histograms of agDD-GFP florescence levels in HeLa^dCas9^ cell line post-S1 washout measured by flow cytometry (left). Histograms of agDD-GFP florescence levels in HeLa^dCas9^ cell line post-S1 washout in the presence of inhibitors measured by flow cytometry BTZ (bortozomib): 1 μM; SAR405 (VPS34 inhibitor): 1 μM (right). **(D)** Histograms of agDD-GFP florescence levels in HeLa^dCas9^ agDD-GFP^LOW^ and agDD-GFP^HIGH^ cells post-S1 washout measured by flow cytometry.

To examine the involvement of proteasomal turnover and to also examine the generality of the apparent agDD-GFP turnover post-washout, we initially created a population HeLa^dCas9^ cells stably expressing agDD-GFP, with an average fluorescence intensity similar to the HEK293T cells (**Fig. 2B and C**). As expected, S1 washout led to the time-dependent accumulation of agDD-GFP puncta, with initial aggregation seen as early as 10 min, as observed by confocal microscopy (**Fig. 2B**). In this cell line, a population of cells with reduced agDD-GFP fluorescence was observed as early as 4 h after S1 washout (**Fig. 2C**, left panel). Importantly, the reduction in fluorescence at this time was completely blocked by addition of the proteasome inhibitor BTZ, but was not blocked by SAR405, an inhibitor of the VSP34 lipid kinase that is required for canonical autophagy (24) (**Fig. 2C**, right panel). Taken together, these results indicate turnover of a population of agDD-GFP aggregates after S1 washout in a manner that depends on the proteasome but not canonical autophagy.

### agDD-GFP aggregate burden and proteasomal turnover

We reasoned that the heterogeneity of fluorescence seen after S1 washout could reflect the levels of agDD-GFP present in individual cells within the population, with higher agDD-GFP levels resulting increased “aggregate burden” and differential degradation potential. To examine this possibility, we first created two additional cell populations using – agDD-GFP^LOW^ and agDD-GFP^HIGH^ – with the fluorescence of the low and high pools representing the lower and higher quartile fluorescence of the parental cell population (**Fig. 2D and SI Appendix, Fig. S1G**). Consistent with our hypothesis, S1 washout of agDD-GFP^LOW^ cells resulted in progressive loss of agDD-GFP fluorescence at 2, 4 and 18 h post-washout (**Fig. 2D**, left panel), with >90% of cells displaying an ∼10-fold decrease in fluorescence intensity at 18 h post-S1 washout (**Fig. 2D**, left panel). In contrast, agDD-GFP^HIGH^ cells displayed essentially no reduction in fluorescence at 2 or 4 h, but with a population of cells with reduced signal with 18 h of S1 washout (**Fig. 2D**, right panel). The pattern observed with agDD-GFP^HIGH^ cells was similar to that seen with the original HEK293T agDD-GFP cells (**Fig. 2A**). As expected, the reduction in fluorescence in agDD-GFP^LOW^ cells with 4 h of S1 washout was fully blocked by BTZ but not by SAR405, consistent with proteasomal turnover and no obvious role for canonical autophagy (**Fig. S1H**).

These data are consistent with aggregate burden contributing to the capacity of cells to promote turnover post-S1 washout. To examine aggregation in agDD-GFP high and low cells directly, we imaged aggregates by confocal microscopy (**Fig. 3A-C**). Thirty min post-S1 washout, agDD-GFP high and low cells contained aggregates of a similar size (<1 μm^2^), although the agDD-GFP^HIGH^ cells had ∼3-fold more aggregates than agDD-GFP^LOW^ cells (**Fig. 3B**). At subsequent time points (2, 4 and 16 h post-S1 washout), the number of aggregates decreased for both agDD-GFP high and low cells (**Fig. 3A**), but the behavior of the aggregates in each cell line was distinct. At 6 h in agDD-GFP^HIGH^ cells, aggregates reach ∼2 μm^2^ in size and the majority of cells retained aggregates, while in agDD-GFP^LOW^ cells, aggregates remained small (<1 μm^2^) and many cells in the population lacked obvious aggregate puncta (**Fig. 3A-C**). Aggregate size continued to increase in agDD-GFP^HIGH^ cells, with residual aggregates reaching ∼4 μm^2^ in size at 16 h, albeit in a small fraction of cells (**Fig. 3A and C**). In contrast, residual aggregates in agDD-GFP^LOW^ cells remained small (<1 μm^2^) (**Fig. 3A and C**). Taken together, these data suggest that aggregate burden results in phenotypically distinct outcomes, with cellular quality control machinery able to remove lower levels of aggregates but higher burden resulting in aggregates that to some extent evade removal in the time scale examined. At longer times of aggregation in agDD-GFP^HIGH^ cells, residual agDD-GFP often adopts a perinuclear localization reminiscent of that seen with aggresomes (25, 26). These results also indicate that the agDD-GFP system can be used to examine the kinetic trajectories of aggregates and their control by cellular quality control machinery.

**Fig 3:**
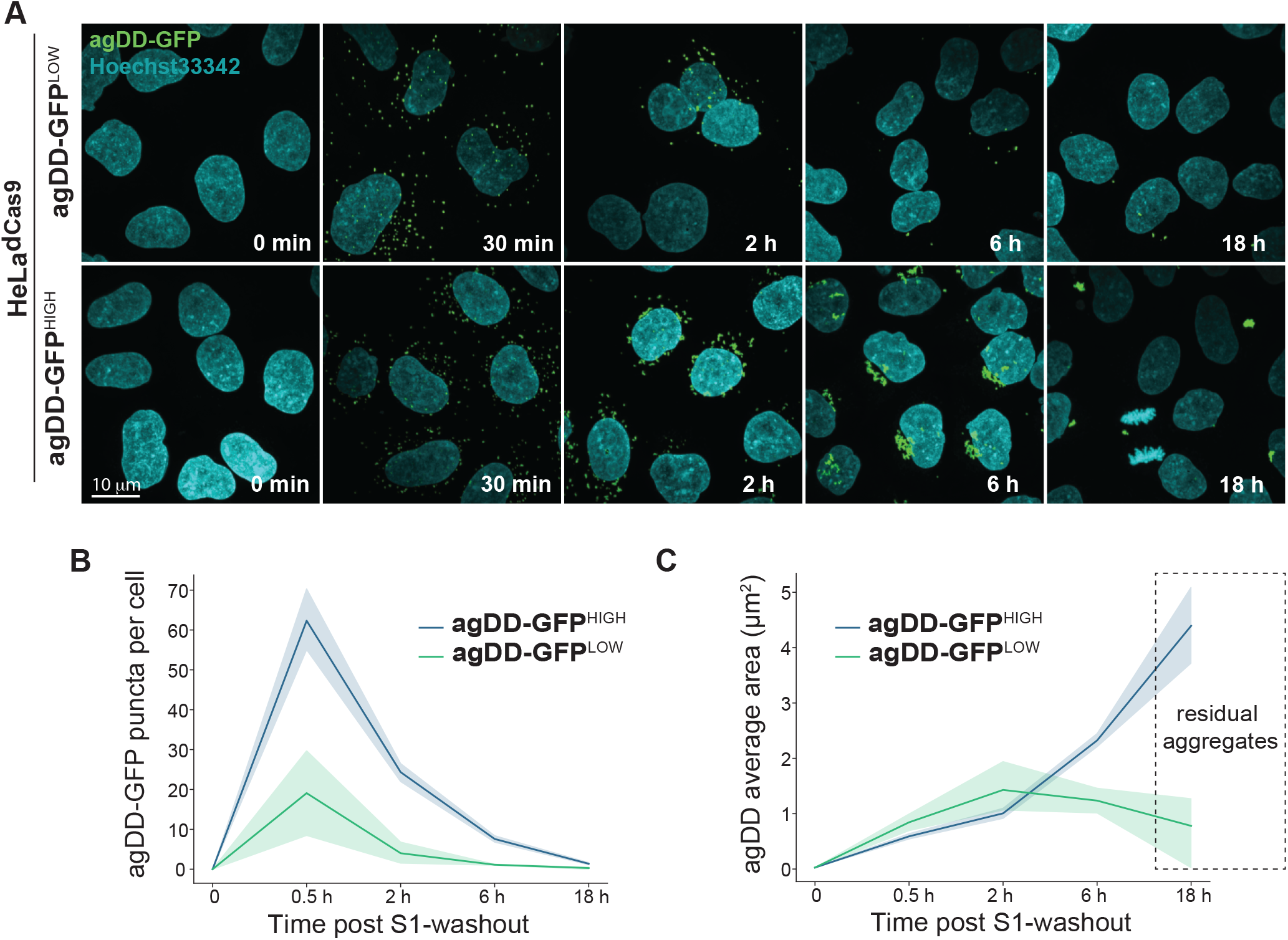
agDD-GFP degradation kinetics is dependent on misfolded protein burden. **(A)** Confocal microscopy max intensity projection of HeLa^dCas9^ agDD-GFP^LOW^ and agDD-GFP^HIGH^ cells at various times post-S1 washout. agDD-GFP signal shown in green, and nucleus in cyan. **(B)** Quantification of agDD-GFP aggregate number per cell at various times post-S1 washout. Line represents mean and shaded area is SD. **(C)** Quantification of average agDD-GFP aggregate size in μm^2^. Line represents mean and shaded area is SD.

### A role for UBE3C but not RPN13 ubiquitylation in proteasomal agDD-GFP turnover

Previous studies revealed that the ubiquitin-proteasome system responds to misfolding of agDD-GFP rapidly (<10 min) to ubiquitylate both agDD-GFP itself, as well as K21 and K34 in a small population (∼1%) of the RPN13 (also called ADRM1) subunit of the proteasome regulatory particle (RP) (13). agDD-GFP aggregation also promotes recruitment of UBE3C to the proteasome, and RPN13 ubiquitylation was suppressed in cells depleted of UBE3C (13). However, it is not clear whether UBE3C E3 activity and/or RPN13 ubiquitylation is involved in cellular turnover of agDD-GFP aggregates. RNP13, through its pleckstrin-like receptor for ubiquitin (Pru)-domain, binds to ubiquitin on degradation substrates (27), and ubiquitylation of K21 and K34 within the Pru-domain could potentially alter RPN13 function in the context of specific substrates. In addition, analogous modifications of RPN13 occur in the context of proteasome inhibition, suggesting involvement in general proteotoxic stress (28). To directly examine the role of UBE3C and related E3s in agDD-GFP turnover, we developed a CRISPRi-based system in parental HeLa^dCas9^ agDD-GFP cells to facilitate analysis of agDD-GFP turnover kinetics by flow cytometry (**Fig. 4A**). Stably expressed CRISPRi sgRNAs for *UBE3A, B, C* in HeLa^dCas9^ agDD-GFP cells strongly suppressed target mRNAs, indicating efficient gene knockdown (**SI Appendix, Fig. S2A**). While depletion of *UBE3A* resulted in significant growth suppression, consistent with cell line sensitivity in DepMap (29), *UBE3B* or *C* depletion resulted in only a small effect on cell proliferation (**SI Appendix, Fig. S2B**). Next, we measured agDD-GFP degradation after 4 h of S1 washout at the population level as revealed by cumulative fraction of cells with GFP fluorescence (**SI Appendix, Fig. S2C**). Only *UBE3C* knockdown had an impact on the cumulative fraction of cells with reduced GFP fluorescence and reversed the fluorescence profile to that seen with cells examined in the presence of S1 (**SI Appendix, Fig. S2C**). This data suggested a primary role for UBE3C in agDD-GFP turnover.

**Fig 4:**
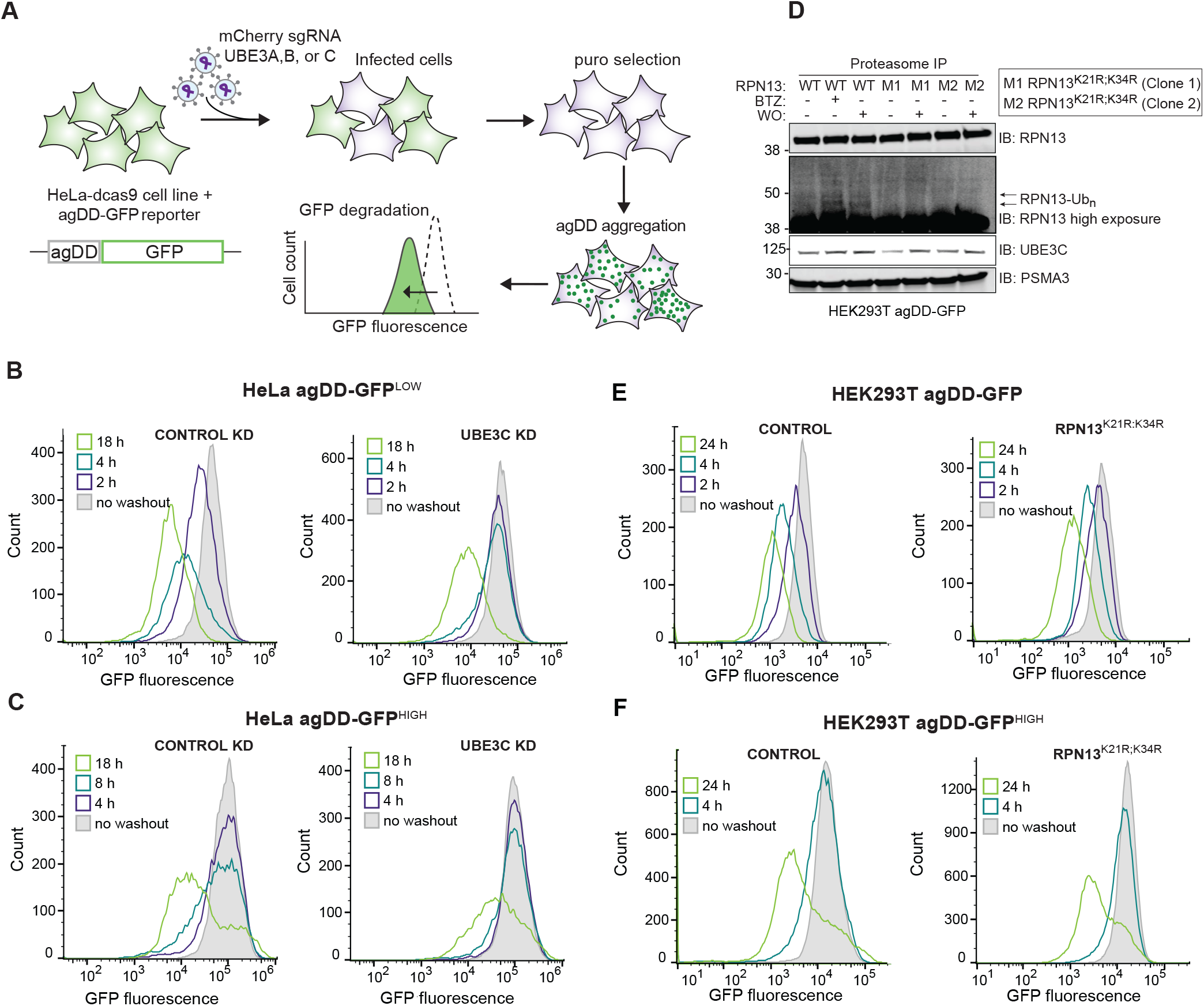
UBE3C is necessary for acute aggregate clearance. **(A)** Workflow for gene knock down in HeLa^dCas9^ cell lines that contain agDD-GFP. First, cells are infected with a guide against the indicated gene (e.g. UBE3C). After selection, S1 is washed out resulting in agDD-GFP aggregation and degradation, which is measured via GFP fluorescence by flow cytometry. **(B)** Histograms of agDD-GFP fluorescence levels in HeLa^dCas9^ agDD-GFP^LOW^ cells post-S1 washout in control cells (left) and UBE3C knockdown (KD) cells (right) measured by flow cytometry. **(C)** Histograms of agDD-GFP fluorescence levels in HeLa^dCas9^ agDD-GFP^HIGH^ post-S1 washout in control cells (left) and UBE3C knockdown cells (right) measured by flow cytometry. **(D)** Western blot of purified proteasomes from HEK293T agDD-GFP and HEK293T agDD-GFP cells with endogenous mutations in RPN13 K21R and K34R post-S1 washout or BTZ treatment. **(E)** Histograms of agDD-GFP fluorescence levels in HEK293T agDD-GFP cells (left) and RPN13^K21R;K34R^ mutations (right) post-S1 washout measured by flow cytometry. **(F)** Histograms of agDD-GFP fluorescence levels in HEK293T agDD-GFP^HIGH^ cells (left) and agDD-GFP^HIGH^ cells with RPN13^K21R;K34R^ mutations (right) post-S1 washout measured by flow cytometry.

To examine the role of UBE3C further, we created HeLa^dCas9^ agDD-GFP low and high cell lines in the context of UBE3C knockdown (**Fig. 4B and C**). In agDD-GFP^LOW^ cells at 2 or 4 h post-S1 washout, UBE3C depletion blocked agDD-GFP turnover (**Fig. 4B**). However, with longer times (18 h), agDD-GFP turnover was still observed (**Fig. 4B**). Similarly, in agDD-GFP^HIGH^ cells, turnover of agDD-GFP was also not fully blocked by UBE3C depletion at 18 h post-S1 washout (**Fig. 4C**). Consistent with these findings and to further validate these results in a different cell line, HEK293T cells edited to delete the *UBE3C* gene (**Fig. S3A and B**) displayed stabilized agDD-GFP at 2 and 4 h post-S1 washout in the context of agDD-GFP^LOW^ and agDD-GFP^INT^ (intermediate) levels (**SI Appendix, Fig. S3C**). However, at 24 h post-S1 washout, a population of cells displayed reduced fluorescence, consistent with agDD-GFP turnover (**SI Appendix, Fig. S3C**). We were unable to successfully generate *UBE3C* knockout HEK293T cells in the context of agDD-GFP^HIGH^. Importantly, the distribution of agDD-GFP fluorescence at 4 h of S1 washout was unchanged upon co-treatment with cycloheximide (**SI Appendix, Fig. S3D**), consistent with the idea that nascent or newly synthesized agDD-GFP is rapidly degraded in the absence of S1 and does not accumulate during the washout period. Taken together, these data indicate a role for UBE3C in agDD-GFP turnover, but with potentially alternative pathways operative over extended periods of aggregation.

The involvement of UBE3C in site-specific RPN13 ubiquitylation in response to agDD-GFP aggregation (13) led us to examine the potential role of RPN13 ubiquitylation by in agDD-GFP turnover. We used CRISPR-Cas9 to introduce homozygous lysine-to-arginine mutations for K21 and K34 in RPN13 in HEK293T cells (**SI Appendix, Fig. S4A**). Importantly, proteomics demonstrated that the abundance of proteasome subunits in GST-UBL-purified 26S complexes from RPN13^K21F;K34R^ cells was largely unchanged when we examined the composition of proteasomes purified from WT cells (**SI Appendix, Fig. S4B, Dataset S1**). Consistent with K21 and K34 representing the major ubiquitylation sites in RPN13 in response to agDD-GFP aggregation (4 h) (13), ubiquitylated forms of RPN13 were observed in WT cells but not in two independent RPN13^K21R;K34R^ clones despite the presence of similar levels of UBE3C in the purified proteasome complexes (**Fig. 4D and SI Appendix, Fig. S4C**). Nevertheless, agDD-GFP turnover in response to S1 washout was unaffected in the RPN13 K->R mutants in both the parental HEK293T agDD-GFP cells and the agDD-GFP^HIGH^ population (**Fig. 4E and F, and SI Appendix, Fig. S4D**). Taken together, these data indicate that the role of UBE3C in agDD-GFP turnover is independent of site-specific ubiquitylation of RNP13, which occurs in response to agDD-GFP aggregation.

### Sustained protein aggregation activates proteotoxic stress transcription factor NRF1

Given the ability of cells to degrade agDD-GFP at extended times after S1 washout with either knock down or deletion of UBE3C, we hypothesized that additional QC pathways may be activated in response to sustained protein aggregation (16). Since our data suggests agDD-GFP degradation is mediated by the proteasome, we hypothesized that proteasome levels may increase over time in response to an influx of misfolded substrates. To test this, we induced agDD-GFP aggregation in the HeLa^dCas9^ agDD-GFP^HIGH^ cells, and measured proteasome subunit levels by qPCR. Within 2 hours of aggregation, we found a ∼1.5-fold increase in *PSMB5* and *PSMD4* mRNAs, which peaked with ∼2-fold increase by 8 hours (**Fig. 5A**). The extent of transcriptional activation was similar to that observed in response to low concentrations of bortezomib (10 nM) measured in parallel (**Fig. 5A**), which has been previously shown to increase proteasome subunit transcription (16).

**Fig 5:**
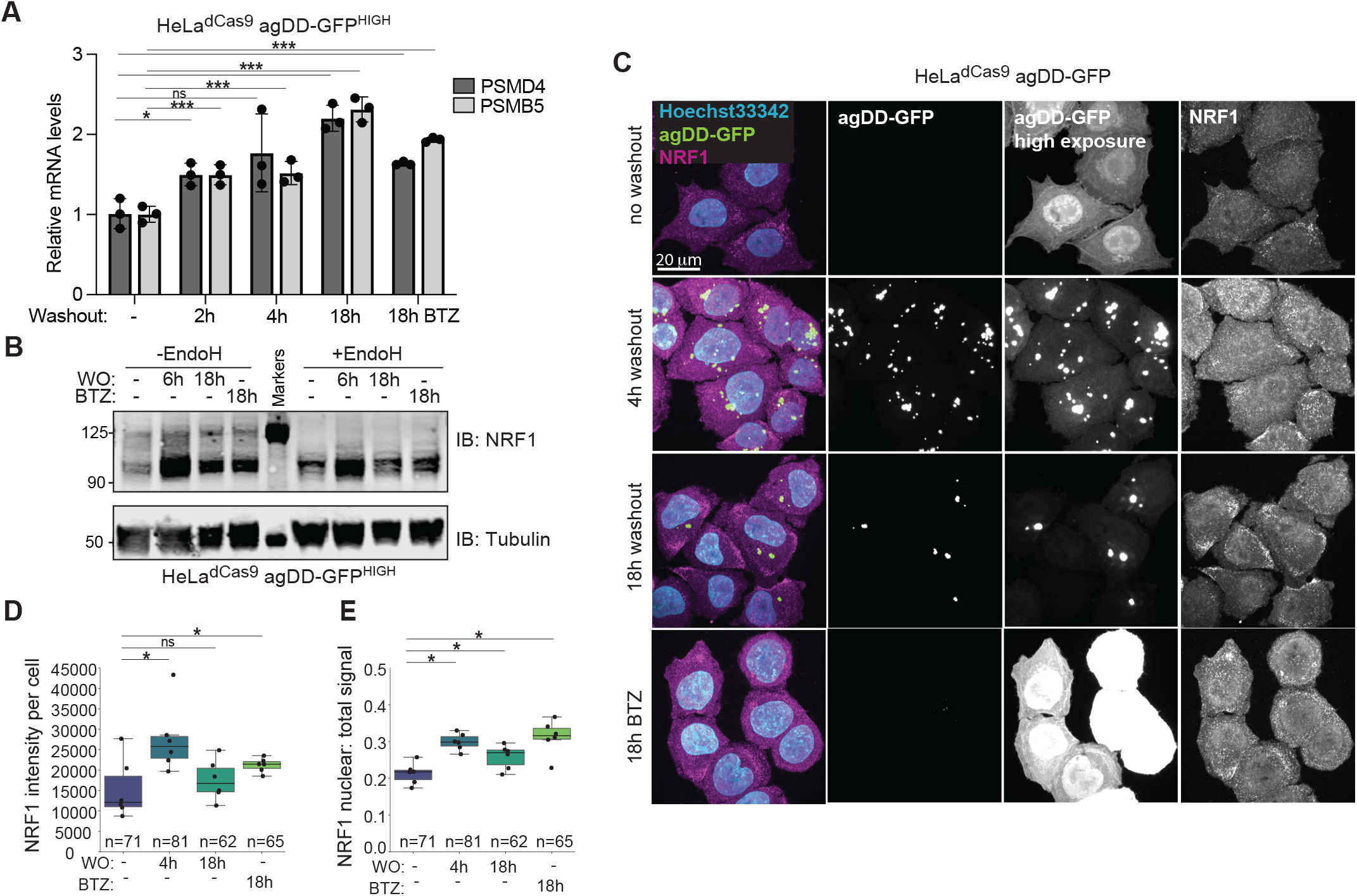
NRF1 is activated by sustained agDD-GFP aggregation. **(A)** Relative mRNA levels for proteasome subunits measured in HeLa^dCas9^agDD-GFP^HIGH^ cells after S1 washout or BTZ treatment (10 nM). * indicates p > 0.05, *** indicates p > 0.01 using a student t-test. **(B)** Western blot of NRF1 protein after S1 washout or BTZ treatment (10 nM). Glycans are removed from NRF1 using EndoH (right). **(C)** Confocal microscopy of HeLa^dCas9^ agDD-GFP cell line post S1 washout or BTZ treatment (10 nM). Nuclei shown in cyan, NRF1 in magenta and agDD-GFP in green. **(D)** Quantification of panel C: NRF1 intensity per cell. * indicates p-value < 0.05 using a student t-test. **(E)** Quantification of panel C: nuclear NRF1 verses total signal. * indicates p-value < 0.05 using a student t-test.

We next reasoned that an increase in proteasome subunit mRNA levels in response to sustained protein aggregation, may be driven by the activation of the NRF1 transcription factor. Although constitutively degraded by the proteasome, upon proteasome inhibition, NRF1 is deglycoslyated, cleaved, and translocated to the nucleus where it increases transcription of proteasome subunits (15–18), as well as proteins involved in autophagy (30). To test if sustained protein aggregation leads to NRF1 stabilization and nuclear translocation, HeLa^dCas9^ agDD-GFP^HIGH^ cell extracts were subjected to α-NRF1 immunoblotting with 6 or 18 h of S1 washout (WO) (**Fig. 5B**). We observed strong induction of NRF1 particularly at 6 h post-S1 washout, as observed with or without treatment of samples with EndoH to collapse glycosylated forms of NRF1 (**Fig. 5B and SI Appendix, Fig. S5A**). At later times (18 h), NRF1 levels were reduced, consistent with reduced aggregate load (**Fig. 5B**). The extent of NRF1 induction at 6 h was greater than that produced with BTZ treatment (18 h). In contrast, the HeLa^dCas9^ agDD-GFP^LOW^ cells that can efficiently degrade the misfolded agDD-GFP within a few hours did not stabilize NRF1 (**SI Appendix, Fig. S5B**). To confirm NRF1 activation and nuclear localization, we examined NRF1 localization by immunofluorescence with and without S1 washout in HeLa^dCas9^ agDD-GFP^HIGH^ cells (**Fig. 5C**). We found that within 4 h of aggregation, endogenous NRF1 levels were increased, and the nuclear ratio of NRF1 was also increased, consistent with aggregation leading to NRF1 stabilization and nuclear translocation (**Fig. 5D and E**). To our knowledge, induction of the NRF1 has not previously been reported in the context of wide-spread protein aggregation in cells. Our results may also explain the previous seemingly counter-intuitive observation that induction of agDD-GFP leads to increased proteasomal activity (13).

### NRF1 increases clearance capacity of cells experiencing widespread protein aggregation

The observation that NRF1 is activated upon widespread agDD-GFP aggregation led us to ask whether the NRF1 pathway contributed to elimination of agDD-GFP. We therefore set up a system in HeLa^dCas9^ agDD-GFP^HIGH^ cells that allowed us to examine gain and loss of NRF1 pathway function. First, we found that CRISPR-mediated knockdown of NRF1 resulted in a statistically significant reduction in the ability of agDD-GFP to be degraded at 18 h post-S1 washout, although the effect was not statistically significant at 8 h washout (**Fig. 6A and SI Appendix, Fig. S6A**). A very similar phenotype was observed with knockdown of DDI2, the protease responsible for release of NRF1 from the ER (**Fig. 6A and SI Appendix, Fig. S6A**). Second, we examined ectopic expression of wild-type NRF1 as well as a constitutively active version of NRF1 (18ND) in which 8 putative N-glycosylation sites have been mutated from asparagine to aspartic acid (31), as initially described in C. elegans (18). We found that both versions of NRF1 were able to increase the extent of agDD-GFP degradation at both 6 and 16 h post-S1 washout (**Fig. 6B and SI Appendix, Fig. S6B**), without altering the basal expression levels of agDD-GFP (**Fig. S6C**). Thus, NRF1 is partially rate-limiting for turnover of agDD-GFP in agDD-GFP^HIGH^ cells.

**Fig 6:**
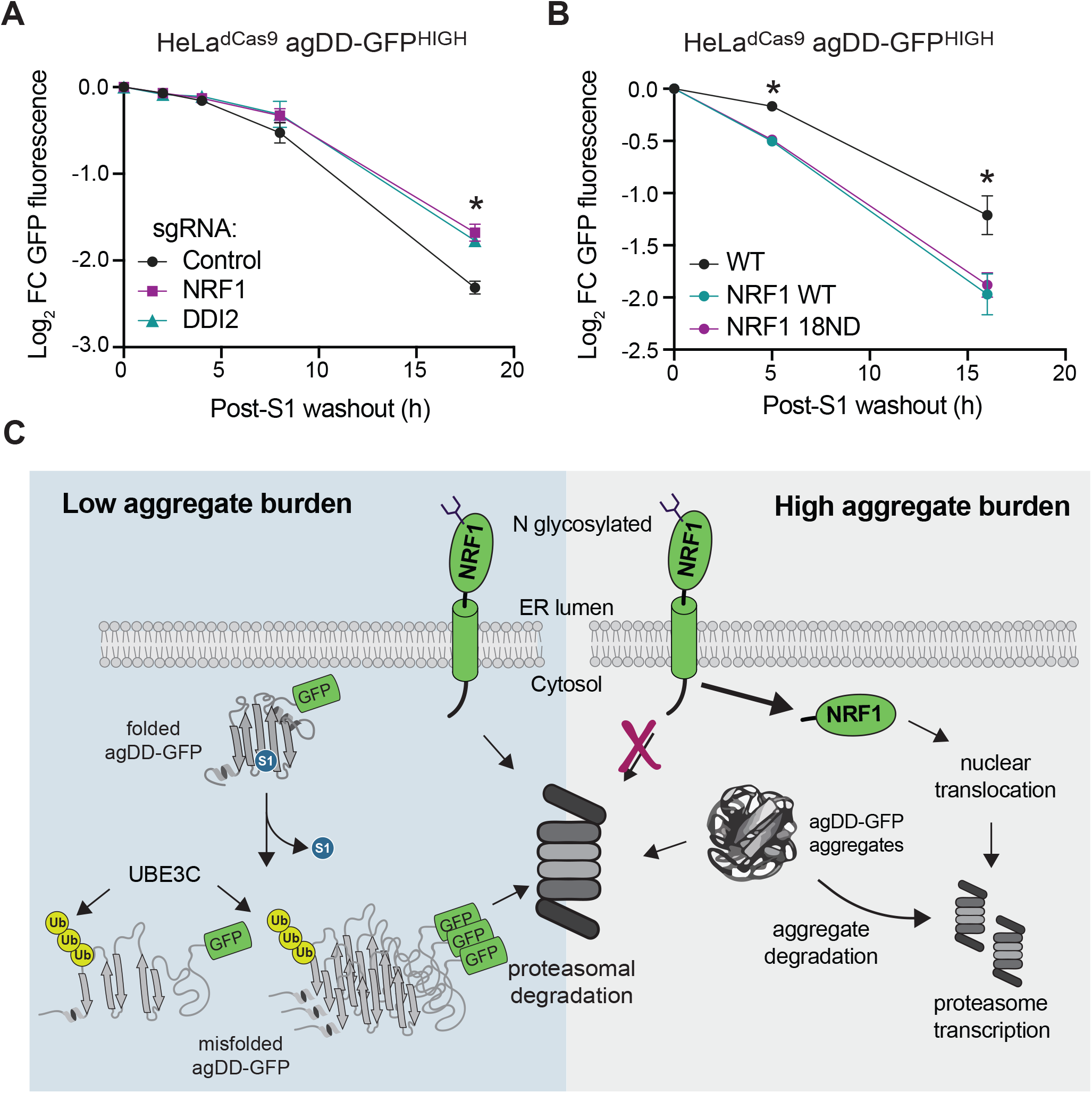
NRF1 modulates aggregation clearance. **(A)** Log_2_ fold change of GFP fluorescence in HeLa^dCas9^ agDD-GFP^HIGH^ cells with NRF1 or DDI2 knockdown post-S1 washout. Error bars show standard deviation. * indicates p-value < 0.05 using a Dunnett’s test. NRF1 vs control p=0.013, DDI2 vs control p= 0.020. **(B)** Log_2_ fold change of GFP fluorescence in HeLa^dCas9^ agDD-GFP^HIGH^ cells that are over-expressing wild-type or 18ND mutant NRF1 post-S1 washout. Error bars show standard deviation. * indicates p-value < 0.05 using a Dunnett’s test. WT NRF1 vs control: 5 h; p=0.00477, 16 h; 0.0345. 18ND NRF1 vs control: 5 h; p= 0.005, 16 h; 0.0482. **(C)** Model of protein quality control pathways that respond to widespread protein aggregation. At low levels of misfolded proteins or small aggregates/oligomers, UBE3C ubiquitylates substrates to promote efficient proteasomal degradation. When levels of misfolded proteins are high, larger inclusions are formed that are increasingly resistant to acute UBE3C-mediated degradation, resulting in activation of NRF1 and downstream proteasome subunit transcription.

## Discussion

Here we merged a temporally controllable aggregate system, with targeted gene depletion or over expression to uncover cellular responses to acute aggregate formation. In essence, this system allows the production of a “pulse” of aggregates, whose size and number are time dependent, to be followed by a “chase” period in which no newly synthesized aggregates are formed. The chase period allows the fate of acutely formed aggregates to be examined kinetically. We find that the burden of misfolded agDD-GFP protein dictates the quality control pathways that are activated to eliminate misfolded proteins and prevent the accumulation of insoluble structures. Our data support a model (**Fig. 6C**) where the initial response to protein misfolding is targeting by UBE3C for proteasomal degradation, likely mediated through substrate ubiquitylation rather than RPN13 K21/K34 ubiquitylation [also thought to be catalyzed by UBE3C (13)]. However, if the aggregate burden is too high for UBE3C and existing pool of proteasomes to efficiently degrade, NRF1 is stabilized and activates a transcriptional profile, including production of additional proteasome subunits, to increase clearance capacity of cells (**Fig. 6C**). This conclusion is supported by our finding that depletion of NRF1 or DDI2 reduces aggregate clearance while ectopic expression of constitutively active forms of NRF1 promote aggregate clearance. The precise mechanisms underlying NRF1 activation by aggregates and whether transcriptional targets beyond proteasome subunits are involved in aggregate degradation remain to be elucidated. Our results may provide an explanation for the previous finding that proteasome activity is increased in cells experiencing agDD-GFP aggregation (13). In addition, the precise mechanism underlying NRF1-dependent proteasome activation/remodeling remains to be determined. In yeast, Hul5 – the ortholog of UBE3C – associates with the proteasome and is important for proteasomal degradation of difficult to degrade proteins (32). In this regard, UBE3C has been reported to function in turnover of difficult to degrade model substrates (33), as well as CFTR during endoplasmic reticulum-associated degradation (34). Further studies are needed to understand the extent to which UBE3C may also function in an analogous capacity to facilitate turnover of other classes of aggregation-prone proteins in the context of NDs. It will also be important to test the extent to which NRF1 and UBE3C may be functionally coupled.

Numerous studies have implicated aggregate degradation by autophagy (aggrephagy) for various types of aggregates (35, 36), although much of this work is based on co-localization of aggregates with autophagy machinery rather than direct demonstration of aggregate flux through the autophagy system. Previous work has also proposed a role for NRF1 in the activation of aggrephagy by upregulation of p62 and GABARAP1L transcription (30). However, in the agDD-GFP system, we do not detect any role for autophagy in clearance, even at higher aggregation burden or at longer time points, as revealed by both deletion of FIP200 and by inhibition of the VPS34 lipid kinase, both of which are required for canonical autophagy. We speculate that although autophagy-specific factors may also be part of the NRF1 transcriptional response, different states of protein misfolding may be more or less accessible by autophagy machinery, with agDD-GFP aggregates being particularly amenable to proteasomal degradation. Recent results indicate that poly-Q aggregates derived from the Htt protein form both amorphous and fibrillar structures over an extended time period of constitutive expression, and autophagosomes are frequently co-sequestered in the amorphous structures based on cryo-ET (21). It remains possible that distinct aggregate forms and/or the kinetics of formation may restrict the various quality control pathways that are capable of removing such aggregates. It will be interesting to examine the potential role of alternative pathways like autophagy under conditions where proteasome or NRF1 function is disabled. Our results suggest that agDD-GFP aggregates, particularly under acute conditions, are able to be acted upon progressively by the proteasomal system to effect virtually complete clearance of aggregates. Further studies are required to understand potential roles of additional quality control factors (e.g. chaperone) which may assist in releasing monomers from aggregates to facilitate ubiquitylation and/or proteasomal degradation.

## Materials and Methods

Detailed procedures related to cell culture, gene editing, biochemical procedures, mass spectrometry, microscopy and flow cytometry are described in *SI Appendix*, Materials and Methods. Statistical analyses were performed using GraphPad Prism (v. 9.4.1) or using python scipy package. All error bars represent standard error of the mean (SEM), or standard deviation (SD) and statistical significance was determined by Dunnett’s multiple comparisons test or by t-test, as specified in the corresponding figure legends.

## Data and Material Availability Statement

The data, code, protocols, and key laboratory materials used and generated in this study are listed in the Key Resource Table alongside their identifiers/catalog numbers and deposited on Zenodo: 10.5281/zenodo.13947535. Proteomics Dataset S1, proteomics parameters READ.ME file, and tabulated data related to this work are deposited on Zenodo: 10.5281/zenodo.13947535. Uncropped blots, raw qPCR data, and flow cytometry data/gating are provided in Zenodo: 10.5281/zenodo.13625620. All images associated with Fig 3A-C and Fig 5C,D are deposited at Zenodo: 10.5281/zenodo.13625614. Proteomic data (.RAW files) are available via ProteomeXchange consortium by the PRIDE partner with identifier PXD055227. Cell lines reported here will be made available upon request with no restrictions, and cell line RRID descriptors are provided in the Key Resources Table. Raw tomograms are deposited at EMDB with the following accession code(s): EMD-51461 (10 min agDD) and EMD-51460 (6 h agDD).

## Acknowledgments

We thank members of the Harper lab for feedback, Ellen Goodall for assistance in protein purification, and Joao Paulo for assistance with proteomics. This work was funded by Aligning Science Across Parkinson’s (ASAP) (J.W.H., R.F.B), NIH RO1 NS083524 (J.W.H.), NIH RO1 AG011085 (J.W.H.), NIH R01 NS042842 (R.R.K.), a Merck-Helen Hay Whitney Foundation Fellowship (K.L.H.), a Helen Hay Whitney Foundation Fellowship (A.M.), and the German Research Foundation (Deutsche Forschungsgemeinschaft) under Germany’s Excellence Strategy (EXC 2067/1-390729940) (R.F.B.). Michael J Fox Foundation administers the grant ASAP-000282 and 024268 on behalf of ASAP and itself. For the purpose of open access, the author has applied for a CC-BY public copyright license to the Author Accepted Manuscript (AAM) version arising from this submission. We acknowledge CITE imaging core (Harvard Medical School) for imaging assistance.

## Author Contributions

This study was conceived by K.L.H., and J.W.H. The experiments and analysis were performed by K.L.H., A.P., E.M.W., T.S., A.M., W.B. and R.F.B. R.R.K. provided critical reagents. K.L.H. and J.W.H. wrote the manuscript with contributions from all authors.

## Competing Interest Statement

J.W.H. is a co-founder of Caraway Therapeutics, a subsidiary of Merck & Co., Inc., Rahway, NJ, USA and is a member of the scientific advisory board for Lyterian Therapeutics. All other authors have no competing interests to declare.

## Notes

### Summary of Updates

This corrects DOIs for Zenodo deposition of data.

