## Supplemental Methods and Supplemental Figures for "Temporal control of acute protein aggregate turnover by UBE3C and NRF1-dependent proteasomal pathways"

*J. Wade Harper

**This PDF file includes:**

Supplemental text: Materials and Methods
Figures S1-S6
Legends for Movies S1 to S2
Legends for Datasets S1
SI References

**Other supporting materials for this manuscript include the following:**

Datasets S1

**Supporting Information Text**

**Materials and Methods**

Information related to all key resources used in this paper can be found at Zenodo (10.5281/zenodo.13947535).

**Antibodies**
Tubulin (Abcam, ab7291; RRID:AB_2241126; dilution 1:1000); IRDye 800CW Goat anti-Rabbit IgG H+L (LI-COR, 925-32211; RRID:AB_2651127; dilution 1:10000); IRDye 680 RD Goat anti-Mouse IgG H+L (LI-COR, 926-680; RRID:AB_10956588; dilution 1:10000); NRF1/NFE2L1 (1:1000; abcam, ab238154); HSP90 (dilution 1:1000; Proteintech 60318; RRID:AB_2881429); PSMA3 (Enzo Life Sciences, #BML-PW8110, RRID:AB_10538395, dilution 1:2000); ADRM1/RPN13 (Cell Signaling Technology, #12019S, RRID:AB_2797798, dilution 1:1000); UBE3C (Bethyl Laboratories, #A304-122A, RRID:AB_2621371, dilution 1:1000 and Invitrogen, PA5-31350, RRID:AB_2548824, dilution 1:1000 ); PCNA (Santa Cruz Biotechnology, sc-56, RRID:AB_628110, dilution 1:2000).

**Chemicals, Peptides, and Proteins**
FluoroBrite DMEM (Thermo Fisher Scientific A, 1896701), Benzonase Nuclease HC (Millipore, 71205-3), Urea (Sigma, Cat#U5378), SDS (Sodium Dodecyl Sulfate) (Bio-Rad, Cat#1610302), Dulbecco’s MEM (DMEM), high glucose, pyruvate (Gibco/Invitrogen, 11995), TCEP (Gold Biotechnology), Puromycin (Gold Biotechnology, P-600-100), Protease inhibitor cocktail (Sigma-Aldrich, P8340), PhosSTOP (Sigma-Aldrich, 4906845001), Trypsin (Promega, V511C), Lys-C (Wako Chemicals, 129-02541), EPPS (Sigma-Aldrich, Cat#E9502), 2-Chloroacetamide (Sigma-Aldrich, C0267), TMTpro 16plex Label Reagent (Thermo Fisher Scientific, Cat#A44520), Hydroxylamine solution (Sigma Cat#438227), Empore™ SPE Disks C18 (3M - Sigma-Aldrich Cat#66883-U), Sep-Pak C18 Cartridge (Waters Cat#WAT054960 and #WAT054925), SOLA HRP SPE Cartridge, 10 mg (Thermo Fisher Scientific, Cat#60109-001), High pH Reversed-Phase Peptide Fractionation Kit (Thermo Fisher Scientific, Cat#84868), Bio-Rad Protein Assay Dye Reagent Concentrate (Bio-Rad,#5000006), and EBSS (Sigma-Aldrich Cat#E3024). Proteasome purification was performed using an established protocol with GST-UBL as an affinity resin (1) (see protocol DOI: 10.1007/978-1-4939-8706-1_18).

**Cell line creation/maintenance and S1 washout**
Protocols for these steps can be found at: DOI: dx.doi.org/10.17504/protocols.io.14egn61xyl5d/v1. HEK293T (human embryonic kidney, fetus, ATCC CRL-3216, RRID: CVCL_0063), HEK293 (human embryonic kidney, fetus, ATCC CRL-1573, RRID: CVCL_0045), $HeLa^{dCas9}$ (2, 3) (cervical carcinoma cell line CCL-2; RRID: CVCL_0030) and $HEK293T^{FIP200-/-}$ (4) cells were grown in Dulbecco’s modified Eagle’s medium (DMEM, high glucose and pyruvate) supplemented with 10% fetal calf serum and maintained in a 5% $CO_2$ incubator at $37^oC$. Cell line authentication was provided by the vendor, and karyotyping (GTG-banded karyotype) of HEK293 cells (from ATCC) was also performed by the cytogenomics core laboratory at Brigham and Women’s Hospital. agDD-GFP (Addgene #78289) was expressed as described previously (5). Cells were maintained at <80% confluency throughout the course of experiments. Cells were maintained in S1 (1 μM, MedChem Express, cat # HY-112210) to maintain agDD-GFP in a soluble folded form. To initiate aggregation, S1-containing mediate was aspirated from the cells, and conditioned media containing 5 μM $FKBP12^{F36V}$ (6) was added to cells.

**CRISPR-Cas9 gene editing**
Protocols for these steps can be found at: DOI: dx.doi.org/10.17504/protocols.io.14egn61xyl5d/v1. CRISPR/Cas9 target sites were selected with the aid of CHOPCHOP (http://chopchop.cbu.uib.no/). Guide RNAs were *in vitro* transcribed using the GeneArt Precision gRNA Synthesis Kit (ThermoFisher Scientific, A29377) and subsequently purified using the RNeasy Mini Kit (Qiagen, 74104). Purified guides were electroporated with purified NLS-Cas9 (Berkeley QB3 MacroLab) at a ratio of 2:1 using the Neon NxT Electroporation System (ThermoFisher Scientific). After a few

days of cell recovery, cells were single cell sorted into 96-well plates by flow cytometry (using a SONY SH800S sorter) to isolate single cells. After 1-2 weeks, single cells that grew into colonies were split into two sets: one for Illumina MiSeq sequencing analysis, the other for cell line propagation. Desired UBE3C deletion clones were confirmed by western blotting. Guide sequence for UBE3C deletion: GAGCTACGAAGACGATGTGGAGG. Guide sequence for RPN13$^{K21R}$ targeting: GGGCGCCTCCAACAAGTACTTGG. Guide sequence for RPN13$^{K34R}$ targeting: GAGTCACGGTGGTCCCCTTCAGG. Ultramer for homology-directed knock-in of RPN13 K21R and K34R: CTCTTTCCAAGCCTGGTGCCAGGCTCTCGGGGCGCCTCCAACA**G**GTACTTGGTG GAGTTTCGGGCGGGAAAGATGTCCCTGA**G**GGGGACCACCGTGACTCCGGATAAGCGGAAA GGGCTGGTGTACATTCAGCAGACGGAC, where the bold nucleotide generates the desired mutation. Generated cell lines were transfected to express agDD-GFP using Addgene #78289, with RRID=CVCL_E3BK for UBE3C and RRID=CVCL_E3BM for RPN13$^{K21R;K34R}$.

**Selective gene depletion (CRISPRi)**

Protocols for these steps can be found at: DOI: dx.doi.org/10.17504/protocols.io.14egn61xyl5d/v1. Single sgRNA expression vectors were individually cloned by annealing complementary synthetic oligonucleotide pairs (Integrated DNA Technologies) for each sgRNA with flanking BstXI and BlpI restriction sites and ligating the resulting double-stranded segment into either BstXI/BlpI-digested pCRISPRi-v2 (Addgene #84832) [marked with a puromycin resistance cassette and BFP as described (7)] or BstXI/BlpI-digested pU6-sgRNA EF1a-puro-t2a-mCherry Addgene #217306) (marked with a puromycin resistance cassette and RFP). Protospacer sequences used for individual evaluation were: UBE3A-GGGCCGCGGCGCAAGACGGG; UBE3B-GCCCGGGTCTGGCAGAACTC; UBE3C-GCACAGCTCGGGCCGCTGCA; NRF1-GAAGCTCCGGCGCCGAGAGTG; and DDI2-GACTCACTGAGCGTGTGTGAG. The resulting sgRNA expression vectors were individually packaged into lentivirus. These sgRNA vectors were transduced into HeLa$^{dCas9}$ cells (RRID: CVCL_E3BH) expressing agDD-GFP (RRID: CVCL_E3BI) at MOI < 1 (20 – 40% infected cells). After selection with puromycin (1μg/ml) for 48 hours, and additional 48 hours of recovery, aggregation time courses were performed.

**Cell lysis and immunoblotting assay**

Protocols for these procedures are provided at: dx.doi.org/10.17504/protocols.io.4r3l226e4l1y/v1. Briefly, cells were cultured in the presence of the corresponding stress to 60-80% confluency in 6-well plates, 10 cm or 15 cm dishes. After removing the media, the cells were washed with DPBS three times. To lyse cell urea buffer (8M urea, 50 mM TRIS 7.5, 150 mM NaCl, containing mammalian protease inhibitor cocktail (Sigma), Phos-STOP, and 20 unit/ml Benzonase (Millipore)) was added directly onto the cells. Cell lysates were collected by cell scrapers and sonicated on ice for 10 seconds at level 5, and lysates were cleared by centrifugation (15000 rpm, 10 min at 4 °C). The concentration of the supernatant was measured by BCA assay. For immunoblotting, the whole cell lysate was denatured by the addition of LDS sample buffer supplemented with 100 mM DTT, followed by boiling at 95 °C for 5 minutes. 10-20 μg of each lysate was loaded onto the 4-20% Tris-Glycine gel (Thermo Fisher Scientific), Bolt 8% Bis-Tris gel, or 4-12% NuPAGE Bis-Tris gel (Thermo Fisher Scientific), followed by SDS-PAGE with Tris-Glycine SDS running buffer (Thermo Fisher Scientific) or MOPS SDS running buffer (Thermo Fisher Scientific), respectively. The proteins were electro-transferred to nitrocellulose membranes and then the total protein was stained using Ponceau (Thermo Fisher Scientific). The membrane was then blocked with LI-COR blocking buffer at room temperature for 1h. Then membranes were incubated with the indicated primary antibodies (4°C, overnight), washed three times with TBST (total 30 min), and further incubated either with fluorescent IRDye 680RD Goat anti-Mouse IgG H+L, or IRDye 800CW Goat anti-Rabbit IgG H+L secondary antibody at (1:10,000) at room temperature for 1h. After thorough wash with TBST for 30 min, near infrared signal was detected using OdysseyCLx imager (LI-COR Bioscience). Uncropped western blots can be found at Zenodo: 10.5281/zenodo.13625620.

**Flow cytometry**

Protocols for these steps can be found at: DOI: dx.doi.org/10.17504/protocols.io.14egn61xyl5d/v1. Corresponding cells were plated onto 24-well, 6-well or 96-well plates one day prior to aggregation. After treatment, cells were lifted using trypsin and quenched with Phenol red free-DMEM. Cells were filtered and analyzed by flow-cytometry (Attune NxT Flow Cytometer (Cat#A28993)-Thermo Fisher Scientific) using the high throughput autosampler (CyKick). The data was processed by FlowJo software (FlowJo; V10.5.2 https://www.flowjo.com) and plotted using GraphPad Prism (Prism; GraphPad, v7&8 https://www.graphpad.com/scientific-software/prism/) and FlowJo. Flow cytometry FCS files can be found at Zenodo: 10.5281/zenodo.13625620.

**Confocal Microscopy**
Image analysis was performed as described: DOI: dx.doi.org/10.17504/protocols.io.q26g717m8gwz/v1. Fixed cells: Cells were plated onto 18 or 22 mm-glass coverslips (No. 1.5, 22x22 mm glass diameter, VWR 48366-227) the day before imaging. After aggregation, cells were fixed using 4% PFA followed by permeabilization with 0.5% Triton-X100. Cells were blocked in 3% BSA for 30 minutes, followed by incubation in primary antibodies for 1h at room temperature. Cells were washed 3 times with DPBS + 0.02% tween-20, followed by incubation in secondary (alexafluor conjugated secondary antibodies) for 1h at room temperature. Coverslips were then washed 3 times with DPBS + 0.02% tween-20 and mounted onto glass slides using mounting media (Vectashield H-1000) and sealed with nail polish. The cells were imaged using a Yokogawa CSU-W1 spinning disk confocal on a Nikon Ti motorized microscope equipped with a Nikon Plan Apo 100x/1.40 N.A objective lens, and Hamamatsu ORCA-Fusion BT CMOS camera. For the analysis, the equal gamma, brightness, and contrast were applied for each image using FiJi software (FiJi ImageJ V.2.0.0 https://imagej.net/Fiji) (8). For quantification, at least 3 separate images. Live cell: Cells were plated onto glass bottom dishes the day before imaging. Before imaging, washout was started to begin aggregation in PhenolRed free dmem. The cells were imaged using a Yokogawa CSU-W1 spinning disk confocal on a Nikon Ti motorized microscope equipped with a Nikon Plan Apo 100x/1.40 N.A objective lens, and Hamamatsu ORCA-Fusion BT CMOS camera, and a live cell chamber with temperature and carbon dioxide control. For the analysis, the equal gamma, brightness, and contrast were applied for each image using FiJi software (FiJi ImageJ V.2.0.0 https://imagej.net/Fiji) (8). Plots and statistics were performed using python packages matplotlib and *python stats*. Image analysis was performed using Fiji (8) and CellProfiler (9, 10) and raw images can be found on Zenodo: 10.5281/zenodo.13625614. Maximum intensity projections of z stacks were converted to tiff images in Fiji, then analyzed in CellProfiler by segmenting nuclei and aggregates and calculating parameters such as number, area and intensity of segmented objects. The output csv file was then loaded into a custom Python script for further analysis and data visualization using pandas, seaborn, numpy, and matplotlib packages. Statistics were performed using stats package. Corresponding p-values were included in figures and figure legends. Python scripts used for analysis and plotting can be found at Zenodo: 10.5281/zenodo.13947535.

**NRF1 overexpression**
Protocols for these steps can be found at: DOI: dx.doi.org/10.17504/protocols.io.14egn61xyl5d/v1. NRF1 (Addgene #181917) and NRF1 18ND (Addgene #181918) (11) overexpression plasmids from were packaged into lentivirus followed by transduction of HeLa$^{dCas9}$ agDD-GFP$^{HIGH}$ cells. Fourty-eight h after infection, cells were selected with puromycin (1$\mu$g/ml) for 48 hours, recovered for an additional 48 hours and then plated for aggregation time course, with S1 washout for the indicated periods of time prior to measurement of GFP levels were measured using flow cytometry. Flow cytometry FCS files can be found at Zenodo: 10.5281/zenodo.13625620.

**qPCR**
Protocols for these steps can be found at: DOI: dx.doi.org/10.17504/protocols.io.14egn61xyl5d/v1. Total RNA was isolated from frozen cell samples using Direct-zol RNA MiniPrep kit. Reverse-transcription was carried using SSIII Reverse-transcriptase (Thermo Fisher Scientific) with oligodT primer (Thermo Fisher Scientific, SO124) in the presence of RNaseIN Recombinant Ribonuclease Inhibitor (Thermo Fisher Scientific). Quantitative PCR (qPCR) was performed with Kappa Sybr Fast qPCR 2x Mix with low ROX (Roche), according to the manufacturer's instructions on Thermo pro quant 7. Experiments were performed in technical triplicates. qPCR primers used: GAPDH: Forward-acgggaagcttgtcatcaat, Reverse-

catcgccccacttgatttt; NRF1 (NFE2L1): Forward-ggaacagcagtggcaagatctc, Reverse-gcaaggctgtagttggtgctca; DDI2: Forward-tgctgaaggaacgcaatccacc, Reverse-cagacgaatcctttcttgctctc; UBE3A: Forward-cgaagaatcactgttctctacagc, Reverse-ggattttccatagcgatcatct; UBE3B: Forward-aagctctgcggaactgtcat, Reverse-cggtgttcgctttcagaac; UBE3C: Forward-ctgccaggatgttcagcttc, Reverse-tttcttcctgagtacgatgtaaaaga; PSMD4: Forward-ctggctaaacgcctcaagaagg, Reverse-cactgtcaccagatgagaaccg; PSMB5: Forward-gtgtcccagaagagccaggaat, Reverse-tcttcaccgtctgggaggcaat. qPCR data can be found at Zenodo: 10.5281/zenodo.13625620.

**Mass spectrometry of purified proteasomes**

The protocol for TMT labeling of purified proteasome can be found at: DOI: dx.doi.org/10.17504/protocols.io.rm7vzjej4lx1/v1. Purified and eluted proteasomes were treated with 10 mM TCEP, 15 mM iodoacetamide, and 10 mM DTT to reduce and alkylate proteins. Methanol-chloroform precipitation was performed on proteins at ratios of methanol to chloroform to water of 4:1:3. Precipitated proteins were washed once with methanol, then spun at 16,000xg, and methanol was removed. Precipitated proteins were resuspended in 100 µL of 200 mM EPPS (pH = 8.0) and digested overnight with endoproteinase Lys-C at 23°C, followed by addition of Trypsin for 6 hours at 37°C, both with a mass ratio of 1:100 protease to total protein. Samples were labeled with 6 µL of TMTpro reagents at a stock concentration of 10 ng/µL and addition of 30 µL of acetonitrile for 1 hour at RT and then quenched with 0.5% vol/vol final concentration of hydroxylamine. Samples were then combined, acidified with 1% formic acid to achieve a pH ~1.5, then subjected to StageTip fractionation, then dried again and resuspended in 1% formic acid 5% acetonitrile for mass spectrometry analysis. Mass spectra were collected on an Orbitrap Fusion Lumos Tribrid mass spectrometer (Thermo Fisher Scientific) using a gradient of acetonitrile from 3 to 13% for the first 83 minutes, then 13 to 28% for the final 5 minutes. An SPS-MS3 method was employed using Real Time Search as previously described (12, 13). Mass spectra were searched using the Comet (14) using a reference human proteome from UniProt (The UniProt Consortium, 2019). A 2% FDR was applied to peptide-spectrum matches (PSM). Protein abundance was quantified from a globally protein-normalized sum of the PSM reporter ion intensity with a signal:noise ratio > 100. Normalization ratios for these proteasome-specific samples were calculated as the average of $\alpha$ and $\beta$ proteasome subunits in each TMT channel and multiplied to each channel. Proteomics data is available on ProteomeXchange with identifier PXD055227.

**Cryo-Electron Tomography (cryo-ET)**

For a description of vitrification, cryo-FIB milling, and cryo-ET imaging methods, see dx.doi.org/10.17504/protocols.io.n92ldm188l5b/v1 and dx.doi.org/10.17504/protocols.io.261ge5pxyg47/v1. Gold 200 mesh R2/1 EM grids (Quantifoil) were coated with approximately 20 nm of carbon in a carbon coater (BAL-TEC MED 020), glow discharged for 45 s in a plasma cleaner, and UV sterilized in a cell culture hood for 30 min. AgDD-sfGFP-expressing HEK293 cells were grown in cell culture medium (DMEM, 10% FBS, 0.5 mM l-glutamine, 1% non-essential AA, 1 µM S1) at 37 °C and 5% $CO_2$. Approximately 25,000 cells were seeded in 35 mm dishes containing four pre-treated EM grids 24 h prior to experiments. Prior to plunge freezing, 5 µM FKBP12$^{F36V}$ (6) was added to titrate intracellular S1 out of the cells. Cells were vitrified at the time points indicated in **Fig. 1A, B** and **Fig. SA-F**.

Cells were vitrified using a home-made manual, gravity-driven plunge-freezer. Before plunge-freezing, cells were treated with 10% glycerol in culture medium as a cryoprotectant for 1 – 5 min (15). EM grids were blotted for 8 s with filter paper from the backside, immediately frozen in liquid ethane/propane and kept at liquid nitrogen temperature.

Grids were mounted on a transfer shuttle designed for a cryo-loading system (16) and loaded into a Quanta 3D FEG dual-beam cryo-FIB/SEM (FEI). Grids were sputtered with platinum (10 mA, 30 s) in a PP3000T loading system (Quorum) to reduce charging effects during electron imaging. Grids were then coated with organometallic platinum as a protective layer for ion beam milling. Grids were imaged with the scanning electron beam operated at 5 kV / 12 pA. Regions of interest (ROI) were thinned down at tilt angles of 18° - 20° with the focused ion beam operated at 30 kV. The beam currents where set to 1 nA at approximately 1 µm distance from the ROI, 500 pA

at 750 nm, 300 pA at 400 nm, 100 pA at 250 nm, 50 pA at 100 nm and 30 pA at 75 nm for fine polishing. If grids were designated for Volta phase plate (VPP) (17) imaging, grids were sputtered once more with platinum (10 mA, 5 s) after milling to increase conductivity of the lamellae.
Samples vitrified at 10 minutes post S1 removal were imaged with a Titan Krios cryo-TEM (Thermo Fisher Scientific) operated at 300 kV, equipped with a FEG, post-column energy filter (Gatan) and VPP. Samples vitrified at 6 hours post S1 removal were imaged using a Polara F30 cryo-TEM (FEI) operated at 300 kV, equipped with a FEG and post-column energy filter (Gatan). In both cases, tilt-series were acquired on K2 Summit direct electron detectors (Gatan) in dose fractionation mode (0.08 frames per second) using SerialEM (18). Tilt-series were taken at -0.5 µm defocus for VPP imaging at 33,000x magnification (pixel size: 4.21 Å) on the Titan Krios and at -9 µm defocus at 22,500x magnification (pixel size: 5.22 Å) on the Polara. The tilt-series were acquired at an angular increment of 2° and typically ranged from -50° to 60°. The total dose was restricted to approximately 120 electrons / Å² per tilt-series.

Frames were aligned and combined using MotionCorr version 1.0 (19). Tilt-series were aligned using patch tracking from the IMOD software package (version 4.8) (19) and reconstructed with weighted back projection. Tomograms were binned four times to a pixel size of 16.84 Å (Titan Krios) or 20.88 Å (Polara) to increase contrast.

Raw tomograms are deposited at EMDB with the following accession code(s): EMD-51461 (10 min agDD) and EMD-51460 (6 h agDD).

### Supplemental Figures

**Fig. S1. Cro-ET of agDD-GFP aggregates and generation of a HeLa$^{dCas9}$ toolkit for examining agDD-GFP aggregate degradation**. (A-C) Examples of tomograms from HEK293 cells with 10 min of agDD-GFP formation after S1 washout. Scale bar, 200 nm. Agg, Aggregate; ER, endoplasmic reticulum; Endo, endosome; Mito, mitochondria; Autolys, autolysosome; Autoph, autophagosome. **(D-F)** Examples of tomograms from HEK293 cells with 6 h of agDD-GFP formation after S1 washout. **(G)** Histogram showing the fluorescence intensity for HeLa$^{dCas9}$ agDD-GFP, HeLa$^{dCas9}$ agDD-GFP$^{HIGH}$, and HeLa$^{dCas9}$ agDD-GFP$^{LOW}$. **(H)** Histograms of agDD-GFP florescence levels in HeLa$^{dCas9}$ agDD-GFP$^{LOW}$ cells (left) and agDD-GFP$^{HIGH}$ cells (right) post-S1 washout measured by flow cytometry, with BTZ (bortezomib): 1 μM; SAR405 (VPS34 inhibitor): 1 μM.

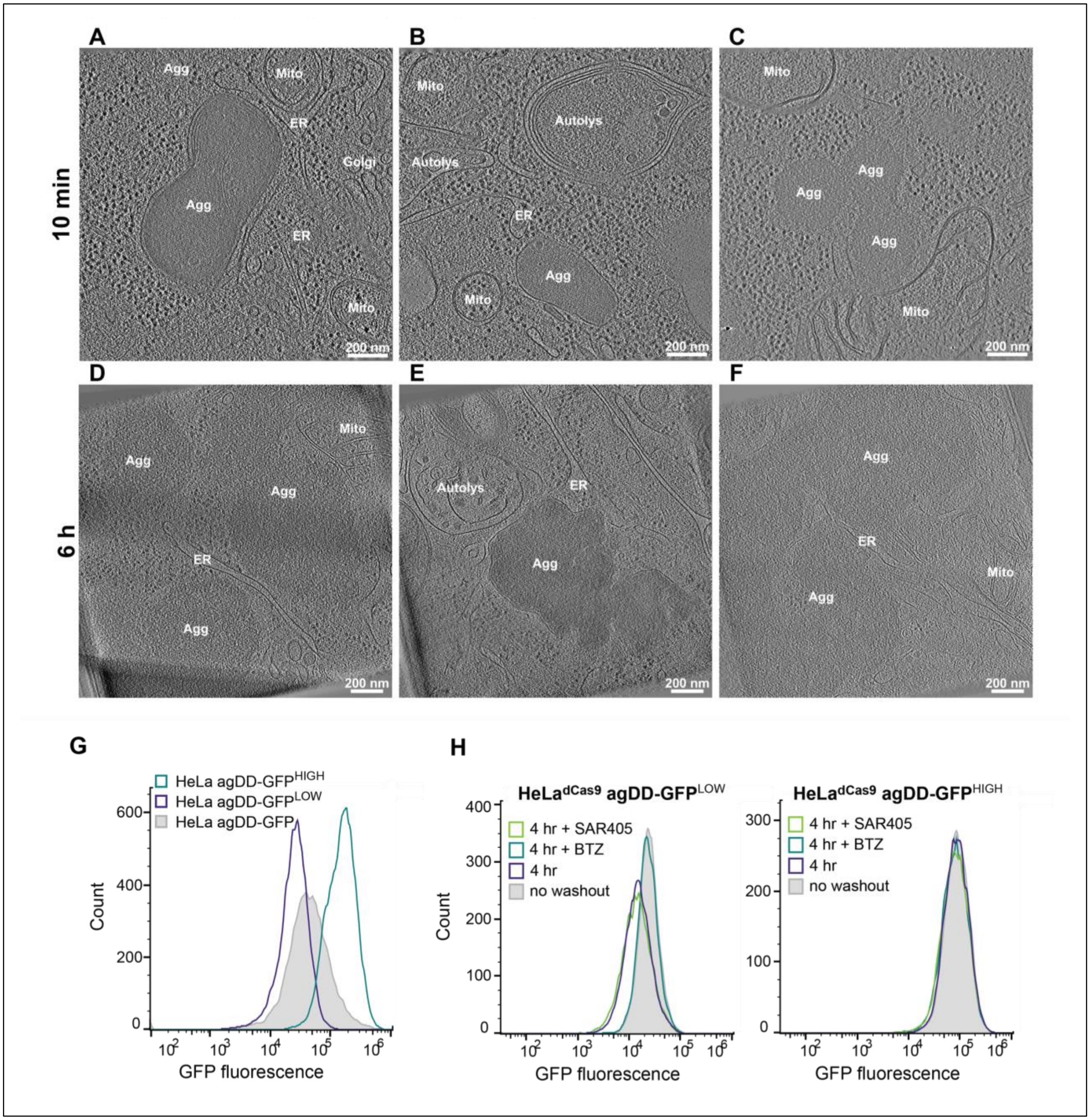

**Fig. S2. UBE3C knockdown impacts agDD-GFP degradation. (A)** mRNA levels for UBE3A, B, and C were measured by qPCR relative to control cells in biological triplicate. Error bars represent SEM. **(B)** Plot of GFP fluorescence at the indicated day post-infection with CRISPRi sgRNAs. **(C)** Plot of GFP fluorescence intensity versus cumulative cell fraction for the indicated cell lines without S1 washout or 4 h post-S1 washout.

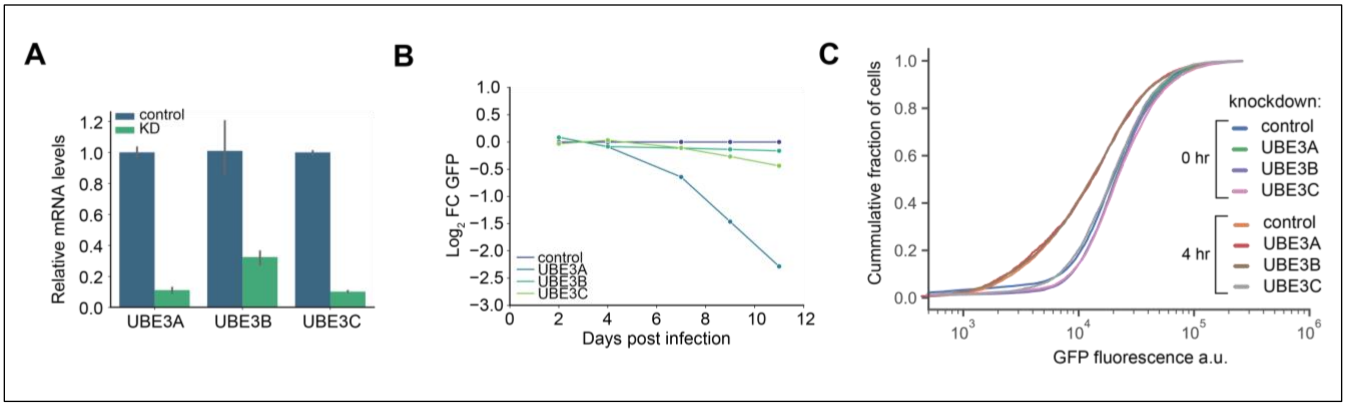

**Fig. S3. UBE3C is required for acute agDD-GFP degradation. (A)** Sequence analysis of two independent clones of UBE3C$^{-/-}$ HEK293T cells, showing in-frame deletions in the indicated alleles. The position of the gRNA is indicated by the yellow shading. **(B)** Immunoblot of wild-type and UBE3C$^{-/-}$ (clone 1) cell extracts using α-UBE3C antibodies, and α-PCNA as a loading control. **(C)** Histograms of agDD-GFP florescence levels in wild-type or UBE3C$^{-/-}$ HEK293T agDD-GFP low, intermediate, or high cells at the indicated times post-S1 washout measured by flow cytometry. **(D)** Histograms of agDD-GFP florescence levels in HeLa$^{dCas9}$ agDD-GFP or agDD-GFP$^{HIGH}$ cells with or without cycloheximide (CHX) with or without S1 washout for 4 h measured by flow cytometry.

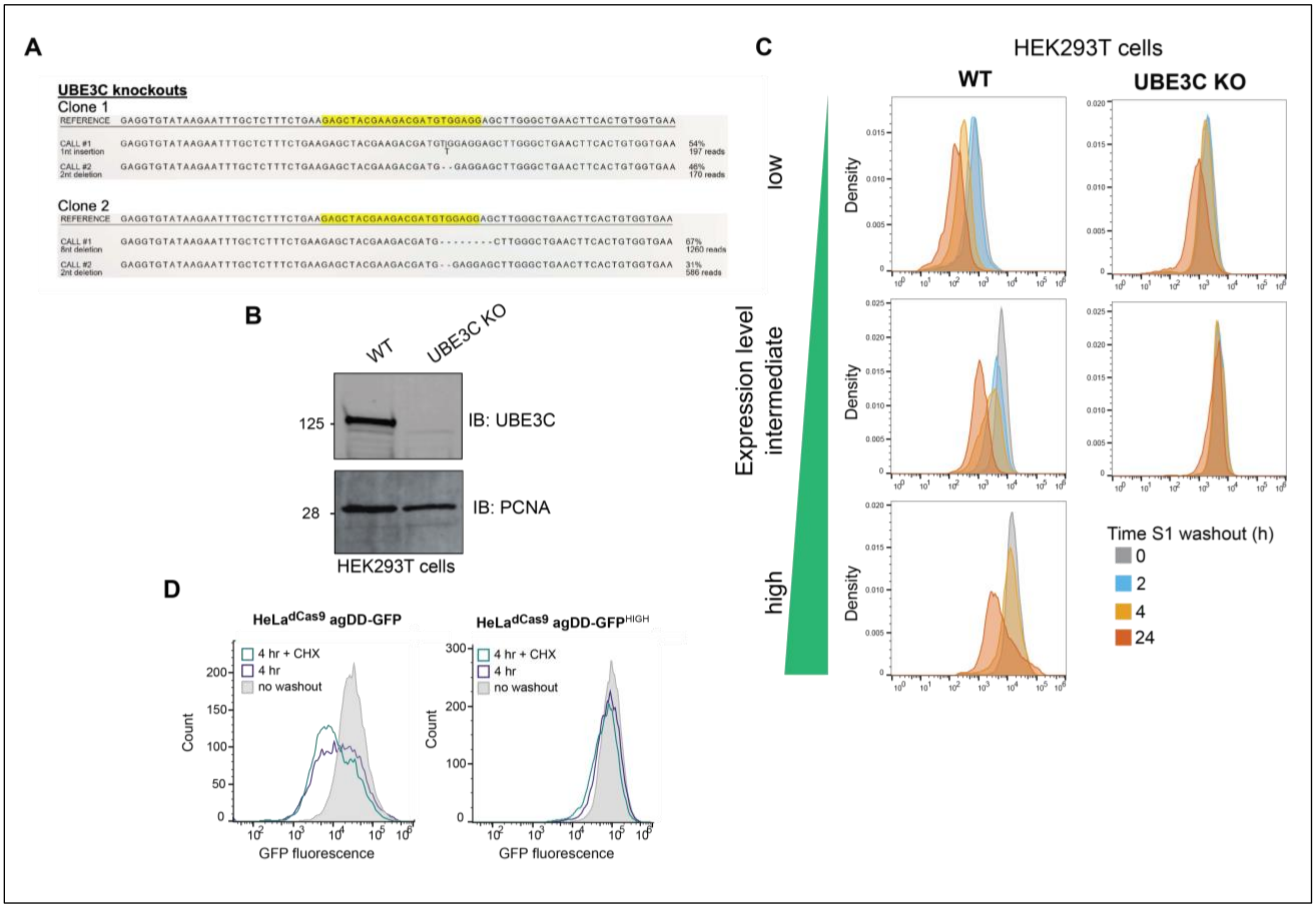

**Fig. S4. RPN13 ubiquitylation on K21 and K34 is not required for agDD-GFP degradation. (A)** Sequence analysis of three independent clones of RPN13$^{K21R;K34R}$ HEK293T cells, showing the desired edits in the indicated alleles. The position of the gRNA is indicated by the yellow shading. **(B)** Tandem mass tagging (TMT)-based proteomic analysis of proteasomes purified from wild-type and RPN13$^{K21R;K34R}$ (Clone 3) cells using GST-UBL affinity resin. **(C)** Immunoblotting of input, flow-through and GST-UBL elution of proteasomes from wild-type and RPN13$^{K21R;K34R}$ HEK293T cells (Clones 1 and 2). Cells were either subjected to S1 washout for 1 h or treated with BTZ for 1 h prior to proteasome isolation. Blots were probed with α-RPN13, α-UBE3C, and α-PSMA3. **(D)** Histograms of agDD-GFP florescence levels in HEK293T agDD-GFP or agDD-GFP$^{HIGH}$ cells with or without RPN13$^{K21R;K34R}$ editing to demonstrate similar levels of agDD-GFP fluorescence measured by flow cytometry.

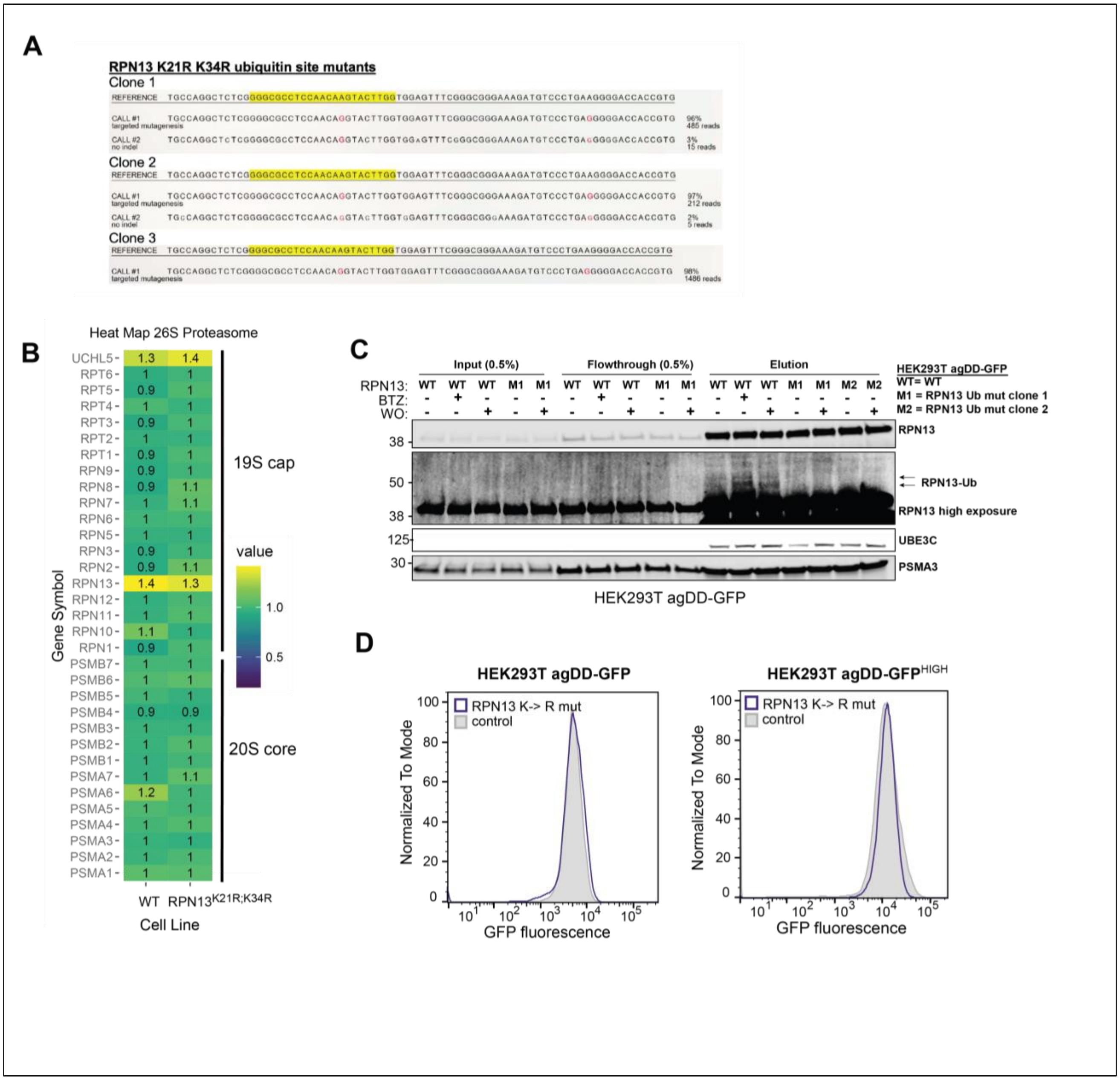

**Fig. S5. NRF1 is activated in response to widespread protein misfolding. (A)** HEK293T agDD-GFP cells were either subjected to S1 washout (WO) or treated BTZ (10 nM) for the indicated time periods. Cell extracts were either subjected to immunoblotting with the indicated antibodies or subjected to EndoH treatment to remove sugar modifications prior to immunoblotting. (B) HeLa$^{dCas9}$ agDD-GFP$^{LOW}$ cells were treated as in panel A for the indicated times followed by immunoblotting of extracts with the indicated antibodies. $\alpha$-Tubulin was used as a loading control. Markers are fluorescent and are therefore detected by LiCOR.

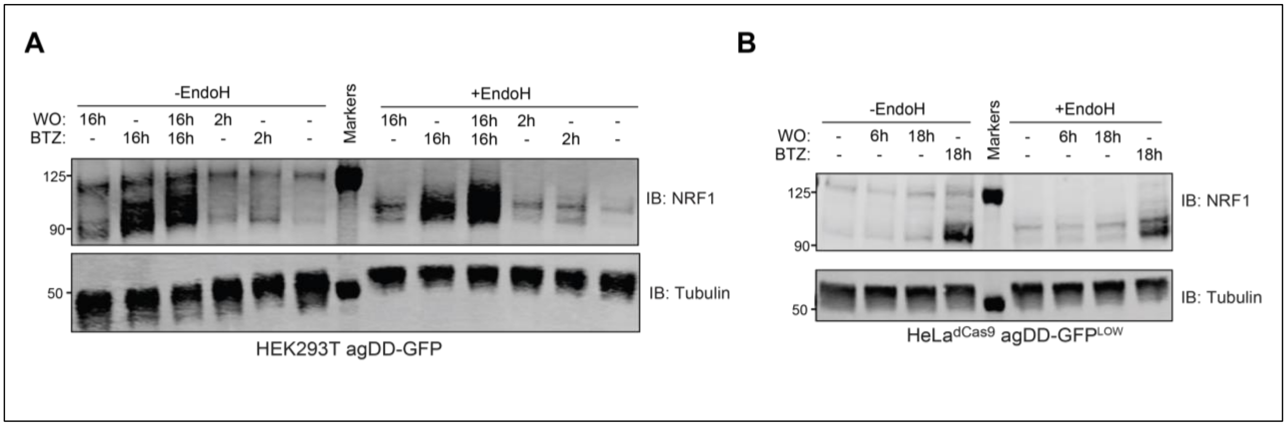

**Fig. S6. Genetic modulation of the NRF1 pathway alters aggregated protein degradation dynamics. (A)** mRNA levels for NRF1 and DDI2 were measured by qPCR relative to control cells. Each point represents the value for individual replicates (n=3 biological replicates). **(B)** Immunoblots of HeLa$^{dCas9}$ agDD-GFP$^{HIGH}$ cells with or without ectopic expression of NRF1 or a constitutively active NRF1 18ND mutant probed with $\alpha$-NRF1 and $\alpha$-HSP90. **(C)** Histograms of agDD-GFP florescence levels in HeLa$^{dCas9}$ agDD-GFP$^{HIGH}$ cells with or without NRF1 or the NRF1 18ND mutant measured by flow cytometry.

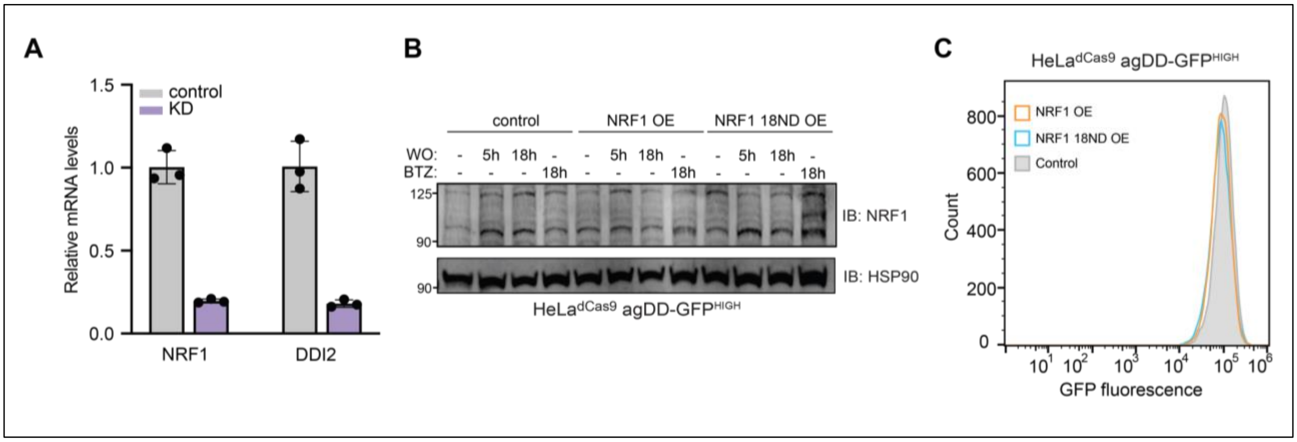

### Legends for Movies

**Movie S1:** Sequential slices back and forth through the representative tomogram of agDD-GFP in HEK293 cells 10 min post-S1 washout in cross-section view. The movie shows amorphous agDD-GFP aggregates often in close proximity with cellular membranes. Related to Fig. 1.

**Movie S2:** Sequential slices back and forth through the representative tomogram of agDD-GFP in HEK293 cells 6 h min post-S1 washout in cross-section view. The movie shows amorphous agDD-GFP aggregates often in close proximity with cellular membranes. Related to Fig. 1.

### Legends for Dataset

**Dataset S1 (separate file).** Proteomic analysis of proteasomes purified from wild-type and RPN13$^{K21R;K34R}$ HEK293T cells. Normalized protein abundance measured by TMT proteomics are indicated. Related to **Fig. S4**. Also deposited at Zenodo (10.5281/zenodo.13947535).
